# 3D-ARM-Gaze: a public dataset of 3D Arm Reaching Movements with Gaze information in virtual reality

**DOI:** 10.1101/2024.01.30.577386

**Authors:** Bianca Lento, Effie Segas, Vincent Leconte, Emilie Doat, Frederic Danion, Renaud Péteri, Jenny Benois-Pineau, Aymar de Rugy

## Abstract

3D-ARM-Gaze is a public dataset designed to provide natural arm movements together with visual and gaze information when reaching objects in a wide reachable space from a precisely controlled, comfortably seated posture. Participants were involved in picking and placing objects in various positions and orientations in a virtual environment, whereby a specific procedure maximized the workspace explored while ensuring a consistent seated posture. The dataset regroups more than 2.5 million samples recorded from 20 healthy participants performing 14 000 single pick-and-place movements (700 per participant). While initially designed to explore novel prosthesis control strategies based on natural eye-hand and arm coordination, this dataset will also be useful to researchers interested in core sensorimotor control, humanoid robotics, human-robot interactions, as well as for the development and testing of associated solutions in gaze-guided computer vision.

## Background & Summary

Every object manipulation starts with a natural arm movement designed to reach and grasp it. Most of the time, this object is recognized based on visual information gathered during gaze-fixation. Understanding eye-hand coordination during reaching movements is a great challenge in human sensorimotor control, with huge applications in motor rehabilitation, humanoid robotics, human-robot interactions, and prosthesis control.

With respect to prosthesis control, recent developments include movement-based control strategies whereby the movement of the prosthetic limb is controlled based on the motion of the residual limb and its natural coordination with the missing joints [1–7]. Although several public datasets exist that include arm movements which could be used to model and exploit this natural coordination [8–10], the overall body postures from which arm movements were produced in those datasets were generally left free or poorly controlled, and none of those datasets were designed to maximize the workspace spanned by participants with their arm (i.e. the set of positions and orientations reachable depending on their morphologies and ranges of motion). Any difference in posture or limit in the workspace explored in those databases as compared to the use case would therefore inevitably translate into malfunctioning control for those posture or workspace area. Furthermore, although 6D pose estimation is a very active area of computer vision [11–14] with applications far beyond prosthesis control (e.g., augmented reality, healthcare or industrial robotics), precise determination of object pose from first person view/egocentric videos with gaze information acquired in natural context is still an open research issue [15]. Here again, several public datasets exist onto which object 6D pose estimation could be trained on and tested [16, 17], but none of those include realistic, egocentric visual information like gaze, obtained in a functionally relevant context (i.e., with natural arm movements that require reaching and displacing objects).

We provide here a dataset designed to overcome these aforementioned limitations. First, the initial body posture from which arm movements were produced was precisely controlled using visual feedback from the trunk and shoulders in a virtual reality (VR) environment. Second, the workspace was maximized by the means of a procedure which initially covers the widest possible workspace (i.e. set by the maximal range of motion of participants), and then subsequently uses a self-organizing map to best represent the space covered by participants (i.e producing natural arm movements within this space) [6]. Third, we recorded head and gaze motion in addition to movements of the trunk, shoulders, and arm joints, thereby collecting the entire kinematic chain between the object the participant is gating (and gazing) at and the hand moved to reach it. This information, combined with the VR technology, enables reconstructing visual data with gaze from which computer vision would need to extract 6D pose of the object of interest in a real-life application. Because the ground truth of the object pose is known by design in VR, and the visual environment could be replayed and manipulated at wish, this dataset can also be used to generate synthetic data consistent with physiological head, gaze, and arm movements, and necessary to develop and test 6D pose estimation algorithms geared toward realistic contexts.

Besides the specific goal of providing information relevant to movement-based prosthesis control [1–7], and in particular those missing in [6] to get closer to a real-life setting, we believe that the present dataset could be useful to a range of broader contexts.

This dataset can be used to revisit several issues lying at the core of human sensorimotor control. First, our 3D recordings of the entire kinematic chain (shoulder, elbow, wrist) are clearly relevant to assess how multiple joints are coordinated in a system exhibiting redundant degrees of freedom (DOF). For instance, this dataset may help to establish whether hand movements are better accounted for by a model in which planning is performed in (intrinsic) joint space or through vectorial coordinates in (extrinsic) task space [18, 19]. One may also be interested in fluctuations of hand and joint kinematics across trials (as we never performed twice the same movement) and assess to what extent a fluctuation from the mean behavior in a given joint might be compensated by a change in another joint [20, 21] so that the resulting hand position is minimally impacted. Because we recorded the maximal range of motion at each individual joint, our dataset is also suitable to investigate to what extent arm configurations can be accounted for by cost functions using postural comfort as a key variable (i.e. maximizing postures away from extreme joint angles [22, 23]). As our dataset was acquired in a large workspace, it can also be used to fuel ongoing debates about the straightness and smoothness of hand movements in 3D space [18, 19, 24–27]. Because our dataset also includes gaze recordings, it can be used to address issues regarding eye-head-arm coordination [28–30]. Finally, how final arm posture may depend on initial arm posture (and initial hand/object position) can also be addressed with the current dataset [19].

On a more applied side, the current dataset offers a benchmark of natural arm movements that can be used in motor rehabilitation contexts (e.g., to generate assistance with upper limb exoskeleton for stroke rehabilitation [31]), or for arm control in humanoid robotics [32–34]. In the context of human-robot interactions, the production of natural arm movements is important as it increases movement predictability, which is known to positively impact our capacity to collaborate with robots [35, 36]. In line with those attempts, the present dataset can be used either to feed methods aiming at reproducing natural arm movement on robotic platforms, or to evaluate the human-likeness of these methods in a wide workspace (see [37] for a review of various methods for human-like arm motion generation).

The following sections describe the methods and content of the dataset, acquired while participants were performing a pick-and-place task illustrated in Supplementary Video 1, with a precisely controlled posture the participants had to return to periodically. The dataset is available on an open repository which also contains basic code for the analyses presented here, as well as a data player allowing both to replay/visualize the data in 3D (Supplementary Video 2), and to generate new video data with various fictitious visual backgrounds and various objects (Supplementary Video 3). This last capability will be particularly attractive to computer vision scientists aiming to develop and test efficient 6D pose estimation algorithms from egocentric vision with gaze. Indeed, being able to increase the variability of objects and backgrounds in the learning phrase can prevent over-fitting.

## Methods

### Participants

The study was conducted on twenty participants (six males and fourteen females), aged 19–44 (mean 25.1; SD 6.46) with normal or corrected-to-normal vision. Participants’ handedness was assessed using the Edinburgh Handedness Inventory (EHI) [38]. All participants were able-bodied, right-handed (mean EHI 81; SD 23) and none of them suffered from any mental or motor disorder that could interfere with the performance of the task. All participants signed an informed consent, and the study was approved by local ethics committee (CCP Est II: n°2019-A02890-57).

### Experimental setup

The participant sat on a stool and wore the headset (ViveTM Pro, HTC Corporation) and five motion trackers (ViveTM Tracker, HTC Corporation), attached to the trunk, the main segments (upper arm, forearm, and hand) of the right arm and the upper arm of the left arm, using straps, as shown in Figure 1a. Each tracker as well as the headset provided measurements of their 3D position and orientation relative to the virtual environment’s reference frame. Two cameras, placed at the corners of the room, received signals from the trackers and the headset at 90 Hz (sampling rate) using SteamVR (Valve Corporation) as middleware. The virtual scene’s contents and interactions with the participant were managed by the Unity simulation engine (Unity Technologies). The virtual environment was scaled to match real-world dimensions: the ground plane was set at the same height of the actual floor and centered in the stool, where the participant was seated.

**Figure 1.**
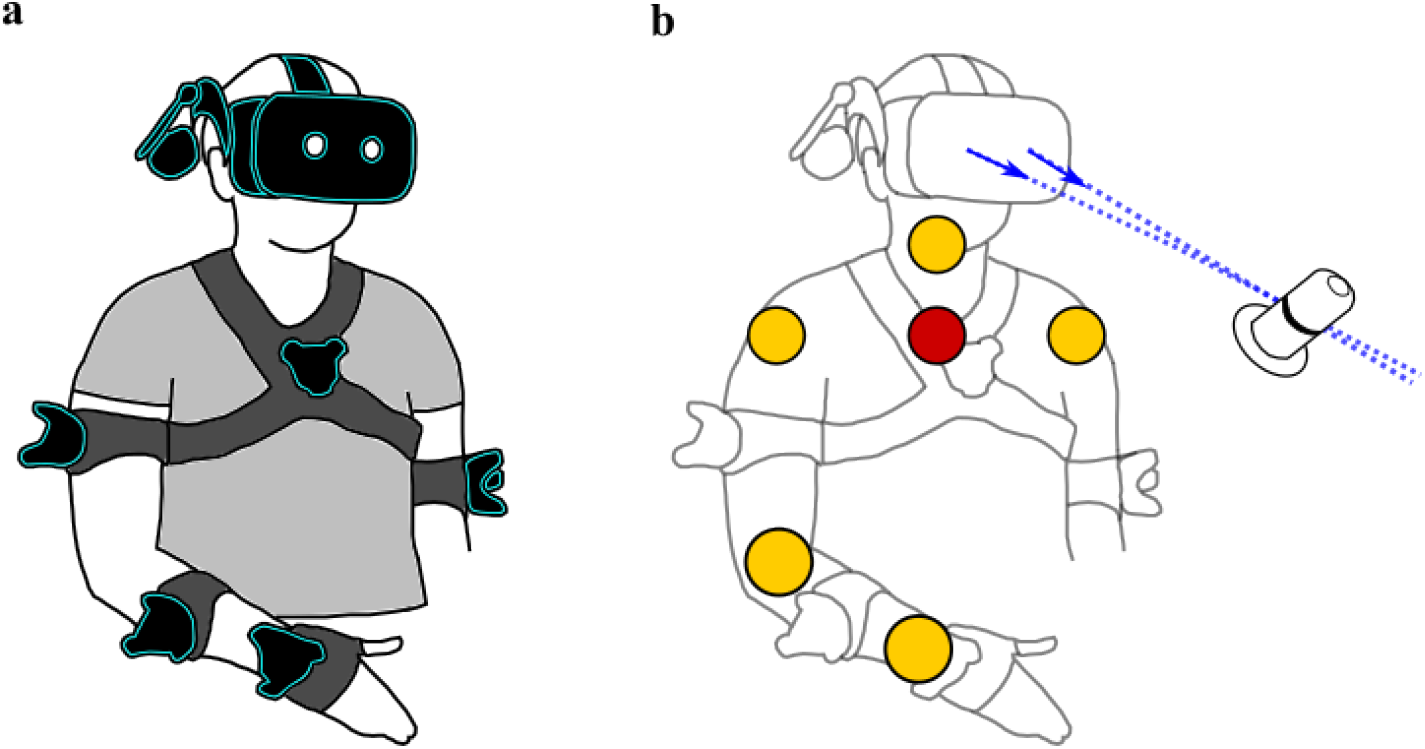
**a**. Participant wearing the VR headset and trackers. **b**. Estimated joint centers’ locations are represented by yellow and red spheres, the pupil axis by blue dashed lines and the direction of the gaze vector of each eye by a blue arrow.

### Calibration

After the participant was equipped with the trackers and the headset, the software for eye tracking was calibrated throughout the method already included in SteamVR [39]. Then, a calibration procedure was carried out to associate the virtual arm with the trackers and therefore, with the participant’s real arm.

The virtual arm, made through the open-source software MakeHuman [40], was composed by three rigid segments (upper arm, forearm, hand) connected by spherical joints at shoulder, elbow and wrist levels. The calibration consisted of three steps: data recording, joint centers estimation, association of virtual and real arm. In the first step, the participant was asked to perform slow movements using all the DOFs of both arms for 15s and then of the head for 10s. Based on these data, the method described in [41] was used to estimate the following joint centers’ locations: left shoulder, neck, right shoulder, elbow, and wrist, shown in Figure 1b as yellow spheres. Additionally, the center of the trunk, represented by a red sphere in Figure 1b, was estimated as the orthogonal projection of the joint center of the neck onto the line connecting right and left shoulder.

Then, the virtual arm (right side) appeared locked in a reference posture with the shoulder placed so that it coincided with the real shoulder of the participant and the virtual arm dimensions resized according to the participant’s ones, computed as the distance between estimated joint centers. In the last step, the participant saw the virtual arm and an arm composed of sticks and spheres, representing his/her real arm. To complete the calibration, the participant had to overlap the arm of sticks and spheres with the virtual one in order to bind them (see [6] for more details).

### Range of motion determination

After the calibration, the headset was temporarily taken off and data were collected with the purpose of computing the range of motion of each DOF (shoulder flexion/extension, shoulder abduction/adduction, upper arm rotation, elbow flexion/extension, forearm rotation, wrist lateral deviation, wrist flexion/extension) of the participant arm (see Supplementary Figure 1). Participants were asked to slowly perform a few repetitions of an elementary movement for each joint with maximal amplitude in each direction. Then, the joint angular limits were estimated as the reached extreme values.

### Task

Data were collected while the participant was engaged in a pick-and-place task involving picking a bottle from one platform (red disk visible at the base of the semi-transparent bottle) and released it into another one, as shown in Figure 2 and in Supplementary Video 1.

**Figure 2.**
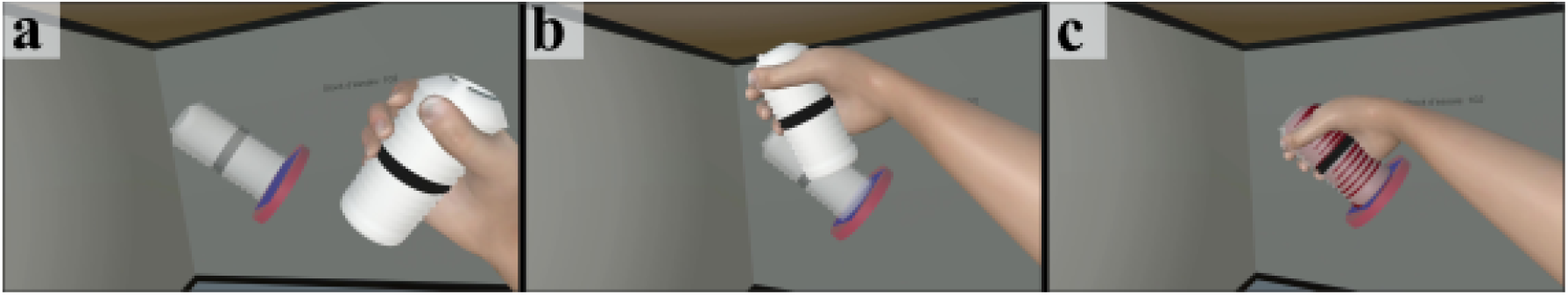
Illustration of the task. After the participant had picked a bottle from a previous platform (*cf* bottle already in the hand), she/he will transport it toward the next platform (**a**-**b**), onto which she/he will release it (**c**). Please note that a shallow bottle has been added on the target platform (**a, b**) to help the participant fitting her/his bottle with the target zone, and that the bottle turned red (**c**) when within the target zone.

One pick-and-place movement was defined as the action from picking the bottle (target) on a platform to releasing on the next platform, or equivalently, from releasing it on a platform to picking it on the next platform. The task did not directly involve the opening and closing of the hand: the participant had to place the virtual hand in the target zone within 2cm of the bottle center and match the virtual hand orientation with that of the bottle within an angular tolerance of 5°. When these two criteria were met, the bottle became red (see Figure 2c), and after 0.6s of continuous red coloring (validation time), the bottle was automatically grasped or released. The participant had 5s to complete each pick-and-place movement: if not achieved within that time, the trial was considered failed and a new target appeared.

### Return to Neutral Posture procedure

In most research focusing on a specific pattern of movement, it is a common procedure to have participants returning to a neutral posture between trials (or movements) in order to maintain recordings consistency and limit drifts or carry over effects between trials [42–44]. As our experiment aimed at collecting data about natural arm movements issued from a comfortable posture, including shoulder and trunk, we designed and applied such a procedure called Return to Neutral Posture (RNP). This procedure was implemented using the display of two sets of three cubes (see Figure 3), each cube corresponding either to a shoulder position (left or right) or to the head position, projected 45cm forward in front of the participant. The green cubes were fixed and represented the target posture the participant had to return to, while the position of the blue cubes was updated according to the actual participant shoulders and head positions. The RNP was achieved by overlapping the blue cubes with the green ones within a given tolerance (angular tolerance 5° and spatial tolerance 2cm for the head and 3cm for the shoulders) for a duration of 0.6s. Supplementary Video 1 shows a representative participant performing the pick-and-place task, with the RNP procedure between each pair of targets (i.e., between each pick-and-place movement).

**Figure 3.**
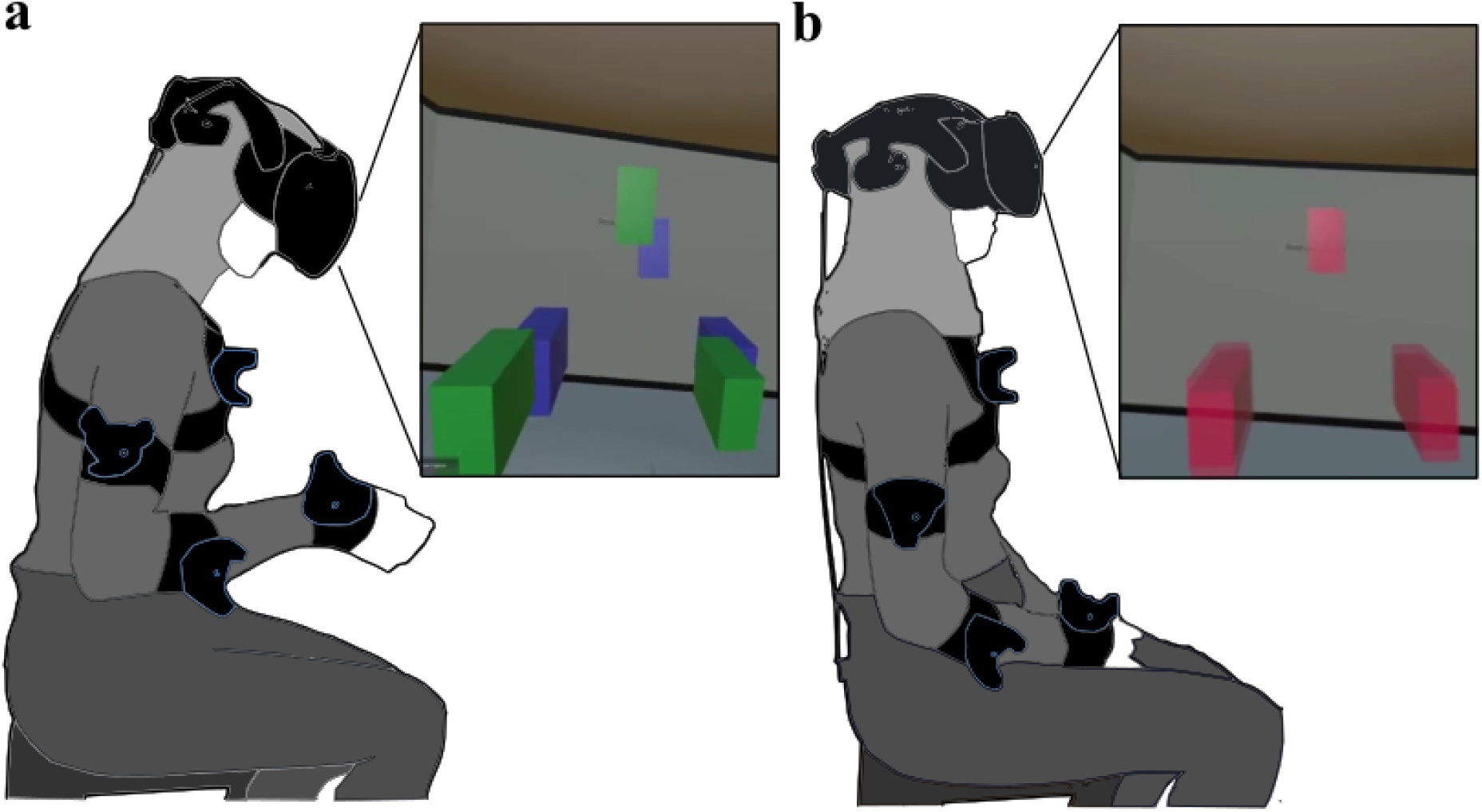
Return to Neutral Posture (RNP) procedure. **a**. After the participant completed a pick-and-place movement, the RNP procedure started and the cubes appeared. The green cubes represented the neutral posture the participant had to return to, while the blue cubes the actual participant shoulders and head positions. **b**. By overlapping the blue cubes with the green cubes within a given tolerance, all cubes turned red to indicate that the participant had successfully achieved the RNP.

### Protocol

The experimental protocol was composed of five phases: Familiarization, Initial Acquisition, RNP familiarization, Test RNP after pauses and Test RNP after target pairs (see Figure 4).

**Figure 4.**
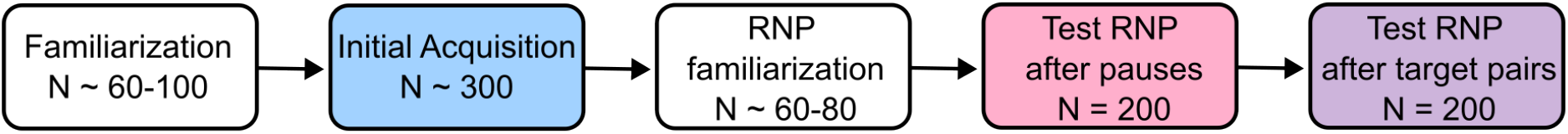
Successive phases of the experimental protocol, with the number of single pick-and-place movements associated with each phase.

Different target sets were generated as explained below for the different phases of the experiment, all being filtered according to the following criteria:

- Exclude targets pointing downward, defined by an angle between the target’s axis and the vertical axis that would exceed 100°.
- Exclude targets too close to the participant’s trunk, defined by a distance between the target’s center and the participant’s frontal plane that would not exceed a third of the participant’s arm length.
- Exclude targets too close to the participant’s legs, defined by a distance between the target’s center and the horizontal plane passing through the participant’s shoulder that would exceed two-thirds of the participant’s arm length.

To generate the first target set, 7-DOF arm angular configurations were drawn at random within the participant ranges of motion following a multivariate uniform probability distribution. Then, forward kinematics was used to compute the target locations (i.e. positions and orientations), which were then filtered as indicated above (see [6] for more details). This target set contained at least 300 targets, subsequently used both in the Familiarization phase and in the Initial Acquisition phase.

The first phase aimed at familiarizing the participant with the virtual environment and the task. For this reason, the time to complete a pick-and-place task was lengthened to 10s. When the participant felt comfortable with the task, after approximately 60-100 targets, the experimenter skipped the remaining targets and proceeded to the next phase.

The second phase, Initial Acquisition (IA), aimed at recording data used to sample more accurately the reachable workspace which is specific to each participant. This was achieved using an unsupervised self-organizing network (more specifically, a Growing Natural Gas (GNG) algorithm [45]), trained on all arm postures produced by each participant during the IA phase, in order to generate a new set of 200 targets that best represent those postures. This step was necessary to circumvent problems inherent to the fact that anatomical joints’ limit are interdependent making some arm configurations impossible, in particular when maximal excursion at multiple joints are involved simultaneously (see [6] for more details). The resulting new set of 200 targets that best represent the reachable workspace of each participant was then used for all next phases.

In the third phase, RNP familiarization, the participants freely choose their neutral posture to return to and got familiar with the RNP procedure, which was introduced every four targets in this familiarization phase. At any time, the participant could ask to reposition the cubes defining that neutral posture, as the goal was to ensure the most natural and comfortable posture. As for the Familiarization phase, when the participant felt comfortable with the neutral posture and the RNP procedure, the experimenter skipped the remaining targets and proceeded to the next phases (typically after 60-80 targets).

Two Test RNP phases were then conducted: one in which the RNP procedure occurred at the beginning of the phase and after each pause introduced every 50 pick-and-place movements (i.e., 3 pauses in a phase of 200 movements), and one in which the RNP procedure occurred after each single pick-and-place movement. This last phase, Test RNP after target pairs (RNP ATP), was designed to prevent any change or drift in baseline posture that might occur from movement to movement. The preceding phase, Test RNP after pauses (RNP AP), aimed at testing whether a RNP procedure introduced only every 50 movements was sufficient to maintain a reliable neutral posture, as this would greatly simplify and accelerate future protocols using this pick-and-place task.

## Data Records

The data recorded during the whole experiment, and the code used for data exploration (see following sections), are available on the open repository Zenodo [46]. The repository also included a Documentation folder with files useful to understand the structure of the dataset and the code provided.

The files are arranged by subject with each subject folder, denominated by the letter “s” and the subject number, containing the following files:

- **“subj_info.json”**: information about the subject (body height and upper limb segment dimensions)
- **“range_of_motion_determination.json”**: data acquired during the range of motion determination
- **“range_of_motion.csv”**: ranges of motion computed from the data acquired during the range of motion determination for each *phase* (Initial Acquisition, Test RNP after pause, Test RNP after target pairs):
  – ***“phase*.json”**: data acquired during the *phase*,
  – **“targets_*phase*.json”**: target set used in the *phase*,
  – **“exclude_targets_*phase*.json”**: lists of targets of the *phase* that should not be considered due to sensor “freeze” or “jump”,
  – **“exclude_times_*phase*.json”**: lists of time intervals of the *phase* containing sensors “freeze” or “jump”,
  – **“exclude_indexs_*phase*.json”**: lists of timestamp indexes of the *phase* containing sensors “freeze” or “jump”.

The files “exclude_targets_phase.json”, “exclude_times_phase.json”, and “exclude_indexs_phase.json” are generated by the code “excludes_generator.py”, which can be found in the folder “code_python” inside “DBAS22_CodeOnline”. This procedure was designed to remove measurement errors associated with motion capture, by applying two filters: one for “freezing” behaviour, defined as a portion lasting at least 0.5s where a sensor’s measurement has not changed value, and one for “jumping” behaviour, when a sensor’s measurement has jumped in position by at least 10cm between two samples. This code generates therefore three lists (one of target numbers, one of time intervals and one of timestamps indexes) of materials that should not be considered when analyzing the data. On average, this filtering process led to the exclusion of 2.6% of targets per participant during each phase.

In the Documentation folder, the file “MainDataExplained” exhaustively describes all the variables recorded. All data is expressed in the world coordinate system if not specified otherwise in the name of the variable.

Two arms information, referred to as the virtual arm and the custom arm, are acquired during the experiment. Both arms are composed of four elements - shoulder, elbow, wrist and hand (end-effector) - for which 3D positions and orientations are recorded. The virtual arm is a 9-DOFs arm directly linked to the sensors’ information, which has been further corrected for errors in humeral rotation. Indeed, soft tissues around the biceps and the triceps are such that the upper arm sensor is not able to follow accurately the humeral rotation. To counter this, we applied the same method as designed in [6] to correct errors in humeral rotation based on additional information from the forearm sensor. Then, the virtual arm was simplified, switching from 9-DOFs to 7-DOFs to obtain the custom arm which matches an ideal anatomical arm description. This was achieved by extracting, from the segment orientations of the 9-DOFs kinematic chain of the virtual arm, the seven joint angles that follow an ideal arm kinematic model, composed of three DOFs at shoulder level, one DOF at elbow level and three DOFs at wrist level (see Supplementary Figure 1, and [6] for further details).

Other data acquired during the experiment include time, position and orientation of the left shoulder, the trunk and the neck, as well as other useful information including the target (position, orientation, type and number) and eyes data (position of the pupils and direction of the gaze for right and left eye).

## Technical Validation

As explained in the Protocol section, the IA phase was designed to generate data useful to sample more accurately the reachable workspace which is specific to each participant, and the two RNP phases were designed to prevent changes or drifts in initial posture that might occur from movement to movement. To assess those, three dependent variables were computed for each participant and phase of the experiment, and subjected to statistical treatment (Friedman tests, followed if significant by post-hoc Conover tests with Bonferroni correction): success rate (the percentage of pick-and-place movements validated in a phase), movement time (the median movement time associated with all validated movements of a phase), and shoulder spread volume (the volume of the ellipsoidal region containing 97% of the shoulder positions recorded while producing all validated movements of a phase, as a proxy of postural stability and body compensation within a phase [2]). Considering the high success rate obtained in all the phases (medians above 96%), the movement time and the shoulder spread volume were calculated on successful pick-and-place movements only. As illustrated in Figure 5, success rates were significantly lower and movement times significantly longer in the IA phase than in the two following RNP phases (IA vs RNP AP vs RNP ATP; n = 20; median success rates of 96.3% vs 98.2% vs 99.4%; Friedman test chi.sq = 24.08, p < 0.001; IA vs RNP AP, p = 0.002; IA vs RNP ATP, p < 0.001; median movement times of 1.61s vs 1.30s vs 1.22s; Friedman test chi.sq = 32.5, p < 0.001; IA vs RNP AP, p < 0.001; IA vs RNP ATP, p < 0.001). This is consistent with the view that some targets, drawn randomly within the range of motion determined individually at each joint of a participant, were difficult to reach, and that the set of targets established on the basis of movements actually produced in the IA phase, and used in the RNP phases, provides a better sample of the reachable space of each participant. The difficulty to reach some targets of the IA phase is further corroborated by four times difference in shoulder spread volume between this condition andthe other RNP phases (IA vs RNP AP vs RNP ATP; n = 20; median shoulder spread volume of 0.76 vs 0.17 vs 0.21; Friedman test chi.sq = 25.2, p < 0.001; IA vs RNP AP, p = 0.001; IA vs RNP ATP, p < 0.001).

**Figure 5.**
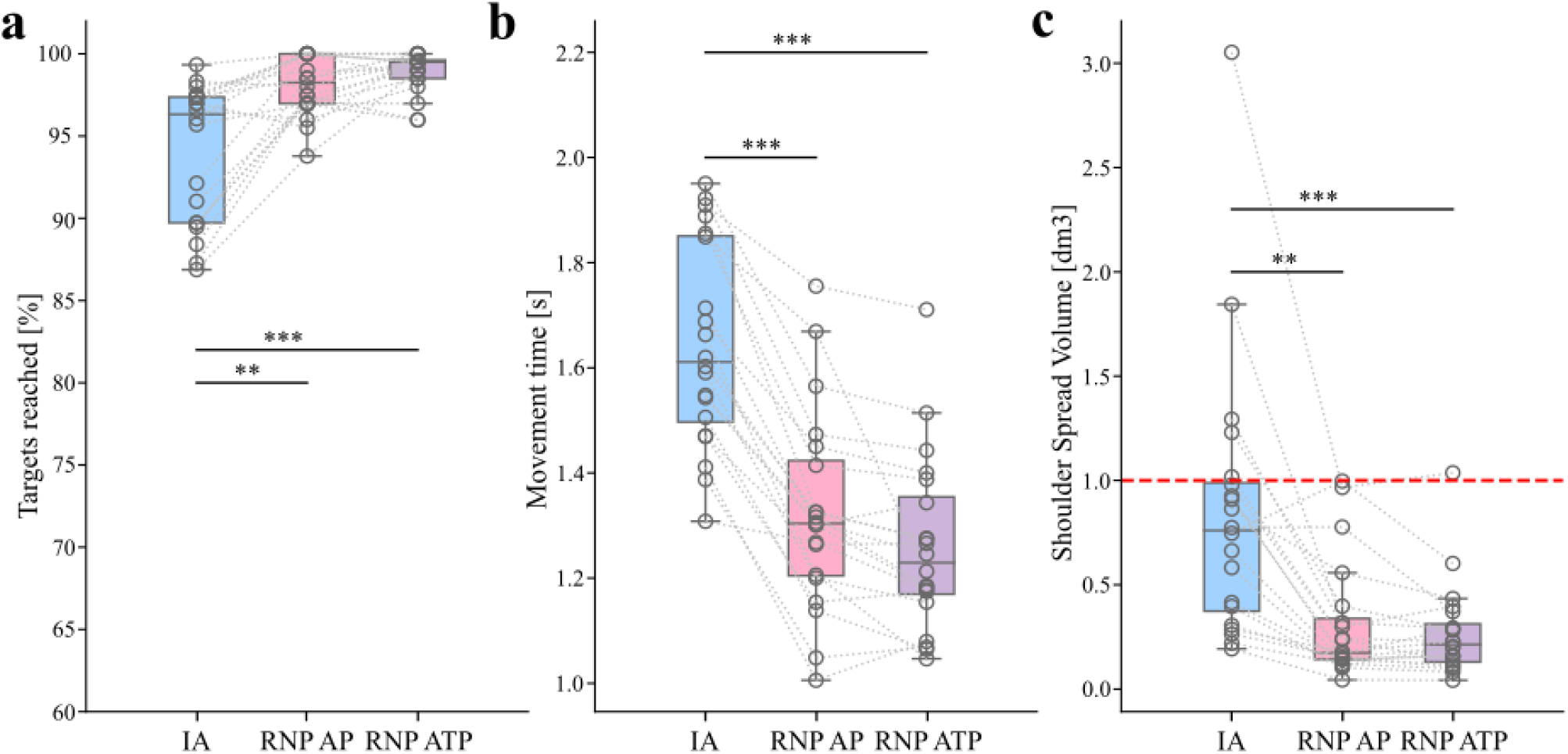
Data analysis results. Individual data are represented by hollow dots and dotted lines. Stars represent significant differences with ** for p < 0.01 and *** for p < 0.001. **a**. Success rate **b**. Movement time. **c**. Shoulder spread volume. The red line represents a volume of 1 dm^3^ (=1 L).

When comparing between the two RNP phases, none of the dependent variables exhibited a significant difference. Pick-and-place movements were therefore performed with similarly high success rates (RNP AP vs RNP ATP; median success rates of 98.2% vs 99.4%; p = 0.997), reliable movement times (RNP AP vs RNP ATP; movement times of 1.30s vs 1.22s; p = 0.366) and led to similar shoulder spread volumes (RNP AP vs RNP ATP; median shoulder spread volume of 0.17 vs 0.21; p = 1), whether the return to a neutral posture procedure was introduced every 50 pick-and-place movements, or between each pick-and-place movements. This lack of difference across RNP phases means that returning to a baseline posture only every 50 pick-and-place movements is sufficient to prevent postural changes or drifts, thereby allowing the release of postural constraints in future protocols using this task.

## Usage Notes

We provide here a brief overview of two methods that we have implemented to analyze and manipulate this dataset: a python code for basic data analyses such as those presented in Figure 5 and 6, and a Unity project called DataPlayer, to playback and visualize previously recorded data, as well as to generate new synthetic data with various visual backgrounds and various objects.

**Figure 6.**
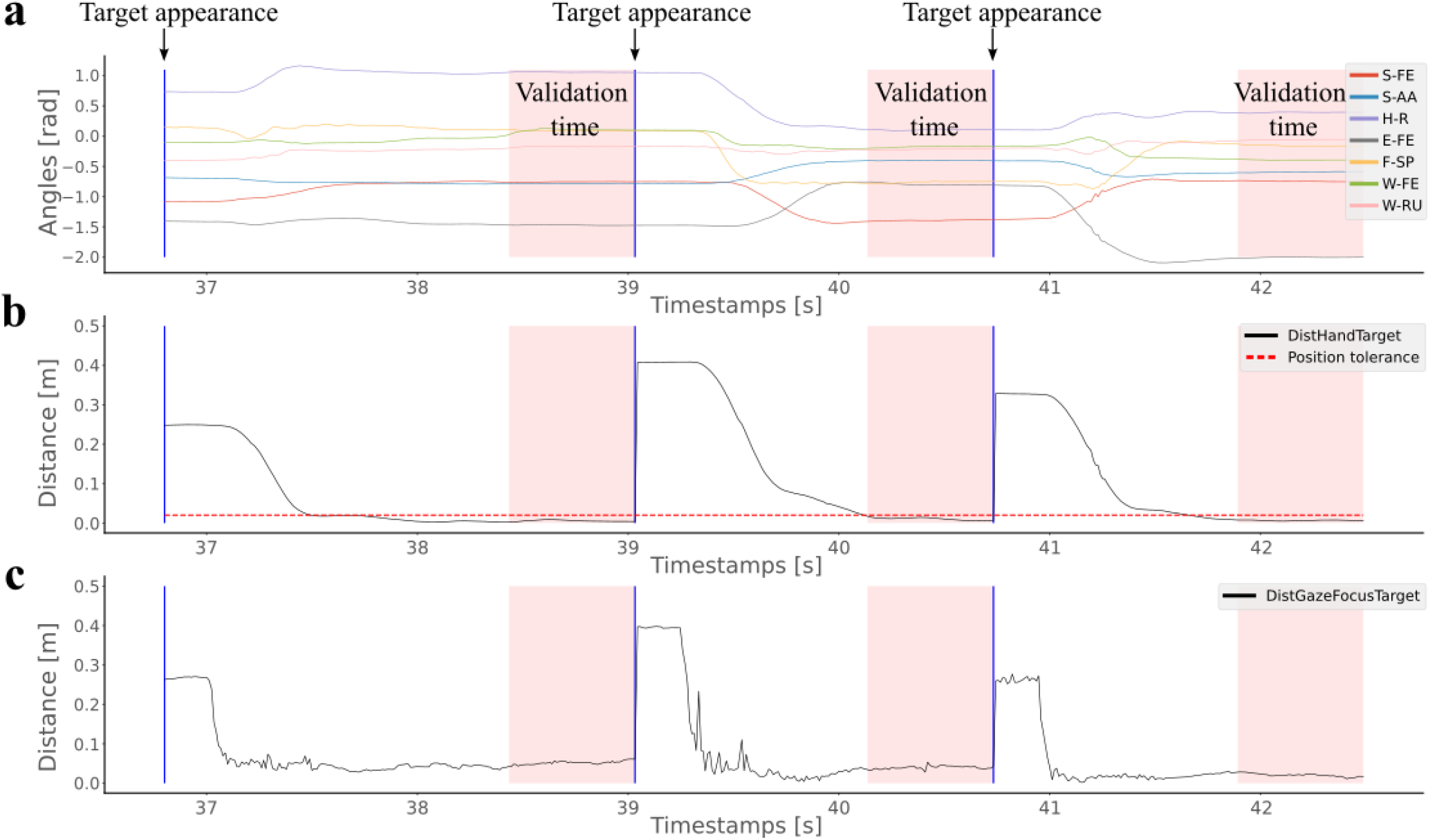
Data collected from participant 8, phase RNP AP, targets 15, 16 and 17. Blue lines represent the target appearance. The red shaded areas represent the time window during which the position and orientation tolerances are met to validate the target (automatic validation after 0.6s of consecutive time within the tolerances). **a**. Time series of joint angles (see Supplementary Figure 1 for a detailed explanation of each angle). **b**. Time series of the 3D distance between the hand and the target. **c**. Time series of the 2D distance (x, y in an egocentric view) between the point of gaze and the target.

### Data analysis

The file “data_analysis.py” in the folder “code_python” contains the code used to analyze the data and generate Figure 5. It includes several useful functions for handling and analyzing the data, such as extracting, filtering the raw data, computing and plotting variables of interest. We also included some basic code to visualize the time series of the angles, of the 3D hand-target distance, and of the 2D gaze-target distance in egocentric views. All of those are displayed in Figure 6, which was generated using this code.

### DataPlayer

We implemented a Unity project, the DataPlayer, allowing to playback the data recorded during the experiment. When selecting a file from the dataset, the DataPlayer reproduces the participant’s arm movements produced during the pick-and-place task, together with gaze data, including the pupil axis and the point of gaze computed with the method described in [47]. Figure 7a shows the interface of the DataPlayer. At the bottom, a control panel allows users to adjust replay speed, trigger pauses, or rewind as needed. In the upper-right corner, some text displays essential information about the data, such as the time, the target number and if the tolerances for target validation are met (tgtRed). In the lower-left corner, a button allows the user to switch between two viewing modes: egocentric and free. In the egocentric view (Figure 7a), the perspective matches that of the participant during the experiment, while in the free mode (Figure 7b), the user can navigate around the scene, explore it from various angles and select a preferred view.

**Figure 7.**
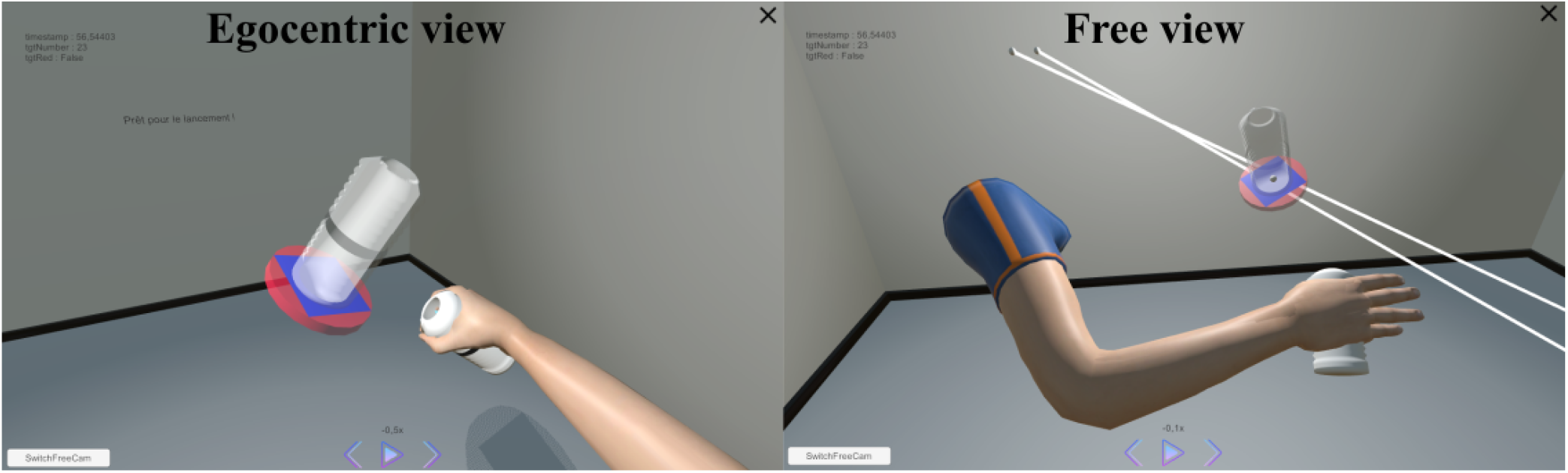
DataPlayer showing an egocentric view (left image) from participant 8 reaching target 23 of the phase RNP AP. DataPlayer showing the same data from a different viewpoint (right image), selected using the free mode. The two white lines represent the pupil axis, and the white sphere the point the participant is looking at.

The DataPlayer being a Unity project, it also allows to modify at wish every aspect of the virtual environment: virtual objects can be added, removed or modified in terms of shape, color and dimension. Customization can be particularly advantageous in applications where the environment needs to reflect a specific context. For instance, the ability to modify objects, attributes, and scenarios provides a fertile ground for computer vision scientists aiming to develop and test robust 6D pose estimation algorithms from egocentric vision with gaze. The project’s malleability allows for simulating diverse real-world scenarios, transforming, for example, the environment into a kitchen setting, complete with culinary items, as shown in Figure 8 and in Supplementary Video 3. Therefore, we have integrated an option in the interface to record and save the newly generated synthetic data, which could be used for training machine learning algorithms to recognize objects in complex environments.

**Figure 8.**
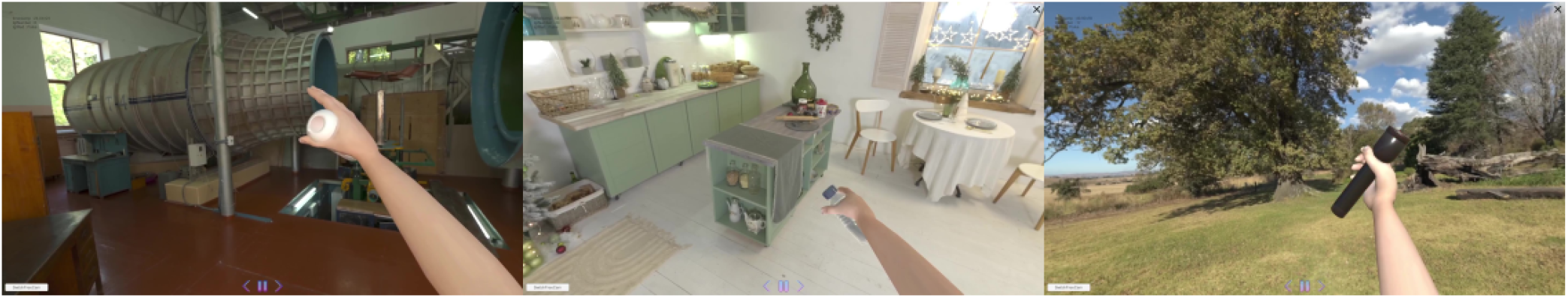
Examples of how the DataPlayer can be customized to combine real data about arm and gaze with various fictitious visual backgrounds and objects.

## Code availability

Data and codes are all available on the open repository Zenodo [46]. This includes the script “data_analysis.py” used for the technical validation and the data analysis described in the Usage Notes, and the DataPlayer which is provided in two versions: a user-friendly stand-alone app which does not require any software installation to replay data as illustrated in Figure 7 and Supplementary Video 2, and a Unity project (folder “code_unity”) which is customizable at wish, as illustrated in Figure 8 and Supplementary Video 3. In the Documentation folder, the file “DataPlayerGuide” contains a tutorial on how to run the project; the file “CodeExplanations” lists and explains all the code scripts related to data analysis; and the document “GuideInstall” provides a tutorial on how to install the environment and the correct packages to run the code.

## Supplementary videos

**Supplementary Video 1 :** https://youtu.be/4ZRYN6ljeCQ

A participant performing the pick-and-place task with the Return to Neutral Posture (RNP) procedure between each pair of pick-and-place movements.

**Supplementary Video 2 :** https://youtu.be/2lSEX_VLK4A

Tutorial on how to launch and use the DataPlayer to replay and visualize data in 3D.

**Supplementary Video 3 :** https://youtu.be/RZhN5IR34_c

Examples of new video data generated with the DataPlayer by combining real data about arm and gaze with various fictitious visual backgrounds and objects.

## Supplementary figure

**Supplementary Figure 1.**
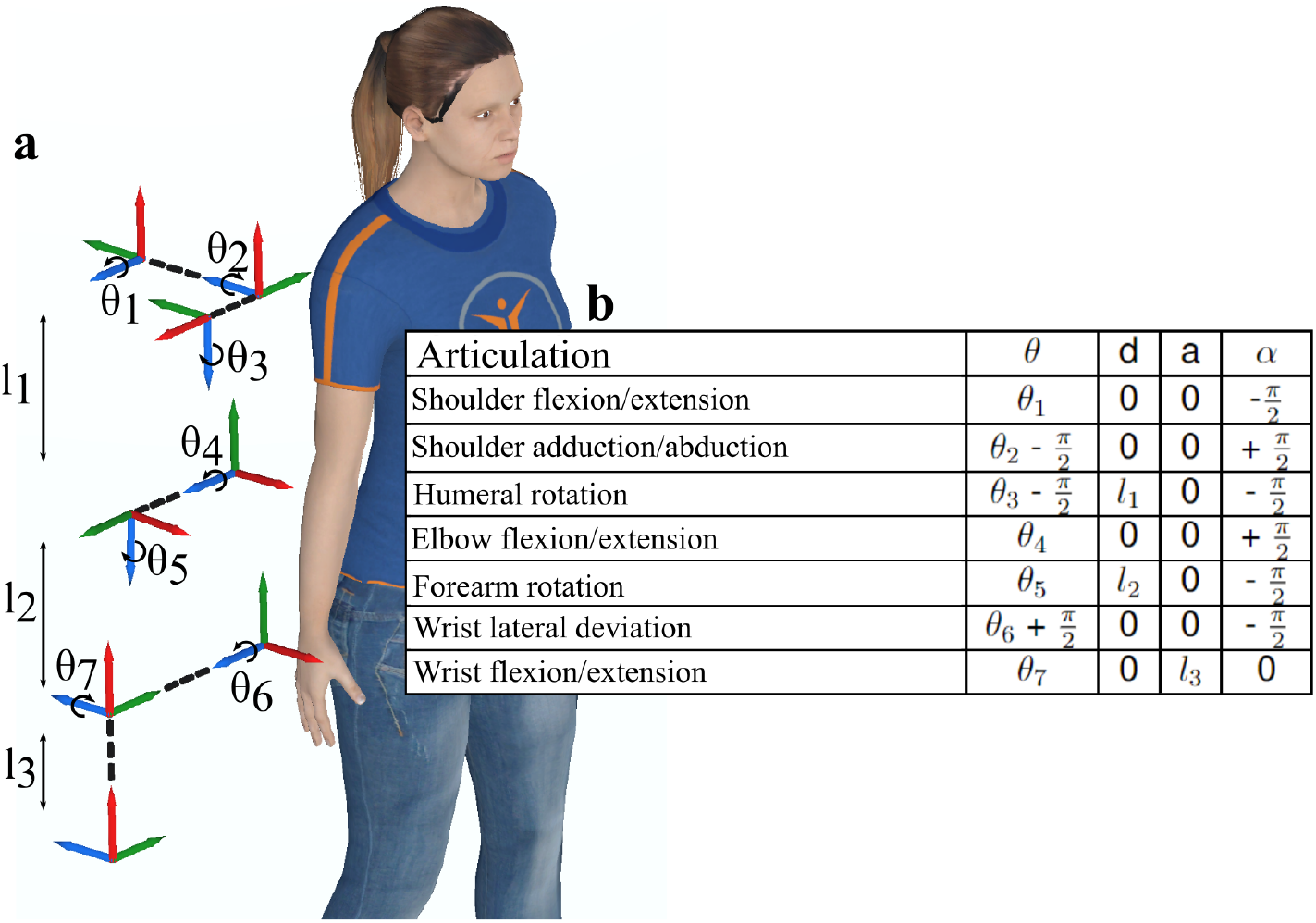
Seven DOF arm model. **a**. Relative positioning of the reference frames. **b**. Denavit-Hartenberg parameters table [48]. (Figure adapted from Figure II.1.5 in [49]).

## Acknowledgements

This work was supported by a PhD grant from the French Direction Générale de l’Armement (DGA) awarded to B.L., and the ANR-PRCE grant I-Wrist (ANR-23-CE19-0031-01) awarded to A.d.R., R.P. and J.B.-P.

## Author contributions statement

A.d.R, J.B.-P and R.P. conceived the study and obtained funding for it. B.L. designed the protocol, performed data acquisition and data processing, organized all materials for the manuscript, and wrote the initial version of the manuscript. A.d.R., E.S. and V.L. participated in the design of the protocol. V.L. contributed to the organization of the materials for the manuscript, including code and data analysis. E.D. performed participants’ inclusion and data acquisition. F.D. contributed to the conception of the study and the writing of the manuscript. All authors reviewed and approved the final version of the manuscript.

